# Hippo Pathway Deregulation Drives Tissue Stiffness and Cancer Stem-like Cells in Lung Adenocarcinoma

**DOI:** 10.1101/390849

**Authors:** Daniela Pankova, Yanyan Jiang, Iolanda Vendrell, Jon N. Buzzelli, Anderson Ryan, Cameron Brown, Eric O’Neill

## Abstract

Lung cancer remains the leading cause of cancer-related death due to poor treatment responses arising from tumor heterogeneity and epigenetic aberrations. Here we show that adverse prognosis associated with epigenetically silenced tumour suppressor RASSF1A is a consequence of increased extracellular matrix, tumour stiffness and metastatic dissemination *in vivo* and *in vitro*. We find that lung cancer cells with methylated RASSF1A display constitutive nuclear YAP1 and expression of prolyl4hydroxylase2 (P4HA2) into extracellular matrix that correlates with increases collagen deposition. Furthermore, we identify an epigenetic axis in tumour cells where elevated ECM impedes the intrinsic suppression of WNT signaling (via TPBG/5T4) triggering b-catenin-YAP1 activation and thus results in a cancer stem-like programming. As key drivers, we identified RASSF1A and P4HA2 mediating the ECM-dependent stemness and metastatic dissemination *in vivo*. Re-expression of RASSF1A or inhibition of P4HA2 activity reverse these effects and increase levels of lung differentiation markers (TTF-1, Mucin5B) *in vivo* and *in vitro*. Our study identifies an epigenetic program to cancer stemness and metastatic progression of lung adenocarcinoma and P4HA2 as potential target for uncoupling ECM signals towards cancer stemness.

## Introduction

Cellular heterogeneity within the tumor microenvironment has been reported as a general property of solid cancers (Hanahan & Weinberg, 2011; Marusyk *et al*, 2014). The population of cells referred to as cancer stem cells (CSCs) exhibit extensive self-renewal abilities, multi-potent differentiation and increased metastatic tumor formation (Ponti *et al*, 2005; Al-Hajj *et al*, 2003; Sales *et al*, 2007; Wang *et al*, 2013). Although CSCs represent <1% of total population of a tumor, they represent a major contributing factor to radio-or conventional chemotherapy resistance and aggressive progression (Kaseb *et al*, 2016; Salcido *et al*, 2010). Dedifferentiation of cancer cells to a more pluripotent-like state results in the appearance of CSCs during malignant tumorigenesis (Codony-Servat *et al*, 2016), however, the mechanism behind this process in solid tumors remains to be elucidated. Recent evidence suggests that interaction with the surrounding microenvironmental niche contributes to the conversion of non-stem cancer cells to CSCs (Gupta *et al*, 2011; Chaffer *et al*, 2011). Both biochemical and physical signals from the tumor microenvironment can modulate properties of CSCs endowing them with the potential to adapt to the emerging cancer niche (Driessens *et al*, 2012; Chen *et al*, 2012; Schepers *et al*, 2012; Schwitalla *et al*, 2013). Mechanical properties provided by extracellular matrix (ECM), e.g. increased levels of collagen, enhanced deposition or crosslinking, have been described to influence CSCs plasticity and trigger cancer stemness in non-stem cancer cells (Ye *et al*, 2014; Wong & Rustgi, 2013). In line with promotion of CSCs, increased matrix stiffness within tumor tissue also correlates with elevated cancer invasion, migration and metastatic spreading (Lane *et al*, 2014; Turner & J. Dalby, 2014; Scadden, 2014; Morrison & Spradling, 2008). Mechanotransduction from the ECM serves as upstream regulator of the Hippo pathway transcription factors YAP/TAZ (Piccolo *et al*, 2014), recently reported as an essential component for formation of lung CSCs (Halder & Johnson, 2011; Tremblay & Camargo, 2012). Nuclear YAP1 is responsible for numerous oncogenic properties of tumor cells and is restricted by the Hippo pathway mediated phosphorylation on serine127 (YAP1-pS127) (Zhao *et al*, 2008; Zanconato *et al*, 2015). RASSF1A is a key regulator of Hippo signaling in humans and loss of expression has been correlated with reduced YAP1-pS127 across multiple clinical cohorts (Vlahov *et al*, 2015). Independently, RASSF1A has been extensively validated as a tumor suppressor in lung cancer where promoter methylation associated gene silencing correlates with poor progression and overall survival (Burbee *et al*, 2001; Lee *et al*, 2001; Neyaz *et al*, 2008; Pallarés *et al*, 2008; Grawenda & O’Neill, 2015). Although RASSF1A has been studied *in vitro*, the precise consequence of loss of RASSF1A expression *in vivo* has been difficult to discern. Gene expression analysis of lung and breast cancers has recently provided insight as, in addition to YAP1 activation, embryonic stem cell signatures are significantly elevated in human tumors lacking RASSF1A (Pefani *et al*, 2016).

To address the precise consequence of RASSF1A loss *in vivo* we constructed H1299 isogenic human lung tumor cells in order to directly assess tumor characteristics in an orthotopic lung tumor model. We found that lung cancer cells with methylated RASSF1A produce prolyl4hydroxylase2 *(P4HA2)* into the extracellular matrix, where is essential for maintaining extracellular matrix-dependent stemness. Concomitantly we found that collagen deposition and tissue stiffness is directly associated with RASSF1A expression *in vitro* and *in vivo*. Moreover, this stiff-ECM displayed strong binding of the CSC marker and suppressor of WNT signaling, trophoblast glycoprotein 5T4 (also known as TPBG or WNT Associated Inhibitory Factor (WAIF1) (Hole & Stern, 1988). We found that 5T4/TPBG fails to suppress WNT in the presence of stiff-ECM and that beta-catenin activation triggers the appearance of a Nanog^+ve^ stem-like cell population in RASSF1A null lung cancer cells. Together, our study provides evidence that widespread clinical prognostic value attributed to RASSF1A epigenetic silencing is due to ECM re-organization associated increase in Nanog^+ve^ stem-like cells and offers new therapeutic opportunities to combat the heterogeneity underlying treatment failures in these cancers.

## Results

### RASSF1A impairs metastases in lung adenocarcinoma

DNA methylation of the CpG island spanning the RASSF1A promoter has been widely appreciated to associate with poor clinical outcome of non-small cell lung cancer (Kim *et al*, 2003; Fischer *et al*, 2007). Unfortunately, to date RASSF1A mRNA levels have not been correlated with epigenetic silencing in NSCLC cohorts, nor have they provided the same prognostic value as methylation. Here we have explored expression levels in The Cancer Genome Atlas (TCGA) and demonstrate that RASSF1A mRNA expression does correlate with a good prognosis for lung adenocarcinomas but not squamous cell carcinomas (Fig 1A and Fig EV1A). To address the physiological role of RASSF1A we constructed isogenic *RASSF1A*-methylated H1299 lung adenocarcinoma cells to stably express RASSF1A or pcDNA3 (control) (Fig 1B). H1299^RASSF1A^ and H1299^con^ cells were orthotopically injected the left lung of mice and examined for tumor formation at day17 or day30 (Fig 1C and Fig EV1B, C). At day 30, 85% of mice (n=5/6) injected with H1299^con^ cells developed clear evidence of primary tumors in left lung and 100% developed contra-lateral metastases in of the right lung (Fig 1C, D, E and Fig EV1C). This process was accompanied by production of liquid oedema around lungs, likely induced by inflammatory response to cancer progression (Movie EV1, 2). Surprisingly, 50% of mice (n=3/6) injected with H1299^RASSF1A^ cells also developed primary tumors similar in size to controls, however, there was a suppression of metastatic events in the both the ipsilateral and contralateral lungs (Fig 1C, D, E and Fig EV1C). This suggests that while RASSF1A can suppress primary tumor growth many, the primary consequence of loss in human tumors may be the failure to suppress metastatic events. Similar results were obtained for primary tumors when sacrificed earlier (day 17) but no metastases were apparent at this time, suggesting that metastases of H1299^con^ are unlikely to be an early event and are a consequence of dissemination from a mature primary tumor (Fig EV1B).

**Figure 1.**
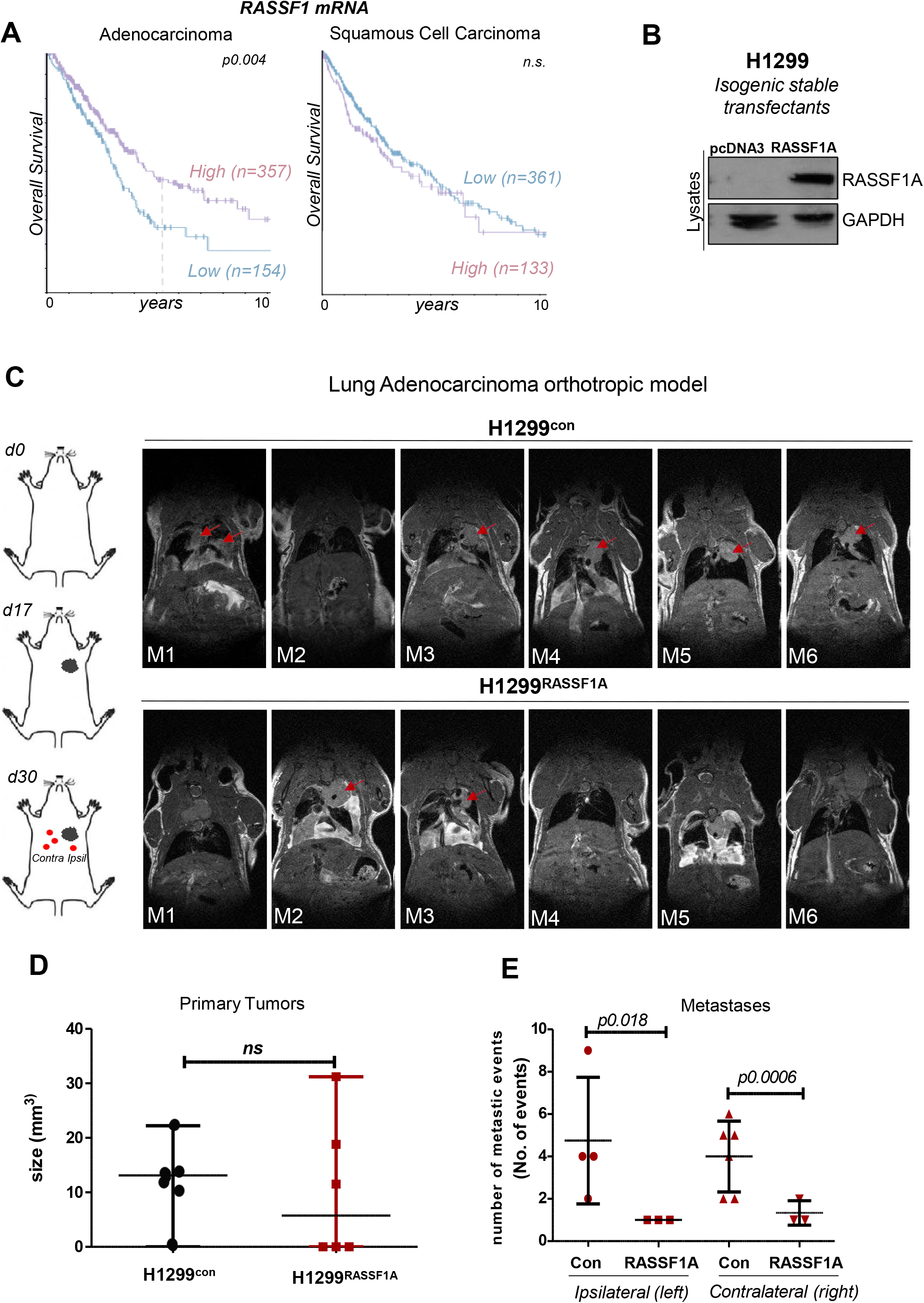
RASSF1A suppresses lung adenocarcinoma metastasis. **A.** Kaplan-Meier curves for lung adenocarcinoma and squamous cell carcinoma patients with high and low expression of RASSF1A mRNA. *The P values were derived from a log-rank test*. **B.** Western blot analysis indicating RASSF1A levels in isogenic H1299 lung adenocarcinoma cell lines. **C.** *Left:* Cartoon of lung adenocarcinoma orthotopic injection. Mice were euthanized to collect the lungs at 17 (*n=4 mice per group*) and 30 days *(n=6 mice per group)* after cell inoculation. *Right:* Longitudinal MR images of a lung tumours in an individual mouse at day 30. Red arrowheads show primary tumours in ipsilateral (*left*) lungs. **D.** Size of primary tumours 30 days after orthotopic injection either with control lentiviral vector (H1299) or with H1299 cells stable expressing RASSF1A protein. **E.** Quantification of metastases in ipsilateral (*left*) and or contralateral *(right)* lungs of each group at day 30. *P values and statistical analyses was performed by using 2-tailed Student’s t-test. Error bars represent S.E.M*.

### Altered mechanical properties of ECM in RASSF1A null tumors

Cancer cell invasion into surrounding tissue is associated with dynamic processes that require ECM and tissue remodeling (Hynes, 2009; Frantz *et al*, 2010; Back *et al*, 2005; Barry-Hamilton *et al*, 2010). In line with reduced metastatic potential, H1299^RASSF1A^ cancer cells had a dramatically reduced ability to invade through 3D-Matrigel matrix compared to H1299^con^ controls (Fig 2A). To further explore processes associated with tissue remodeling during invasion, cells were assessed for their ability to contract a 3D-collagen matrix *in vitro*. After four days, extensive matrix remodeling and contraction of collagen was apparent in the presence of control cancer cells, whereas collagen surrounding H1299^RASSF1A^ cells remained unaltered (Fig 2B). Moreover, substantially greater collagen deposition and organization was apparent around H1299^con^ tumors *in vivo* as visualized by second harmonic generation (SHG) (Fig 2C). YAP is a mechanical sensor of ECM stiffness that facilities growth and expansion of stem cells and tumor cells and intriguingly, it is regulated by RASSF1A. Nuclear localization of YAP occurs on stiff matrix and indeed we observed this in H1299^con^ cells in threedimensional 2% collagen, however in line with its role in Hippo pathway activation, re-expression of RASSF1A prevented nuclear accumulation (Fig 2D). To determine if this was cell intrinsic or due to the “specific” ECM, we extracted ECM from H1299^con^ or H1299^RASSF1A^ cells and used these as a substrate for naïve H1299^RASSF1A^ cancer cells (where YAP is more cytoplasmic due to activated Hippo pathway). YAP transduced to the nucleus in H1299^RASSF1A^ cancer cells grown on the ECM^con^ but not on ECM^RASSF1A^, suggesting that the H1299 control cancer cells with epigenetically silenced RASSF1A produce stiffer ECM that is key for cancer cell re-programming (Fig 2E). Taken together, the lack of 3D-collagen contractility and producing less stiff ECM of H1299^RASSF1A^ cancer cells is consistent with reduction of invasion *in vitro* and *in vivo*, and decreased potential to form primary tumors and metastases.

**Figure 2.**
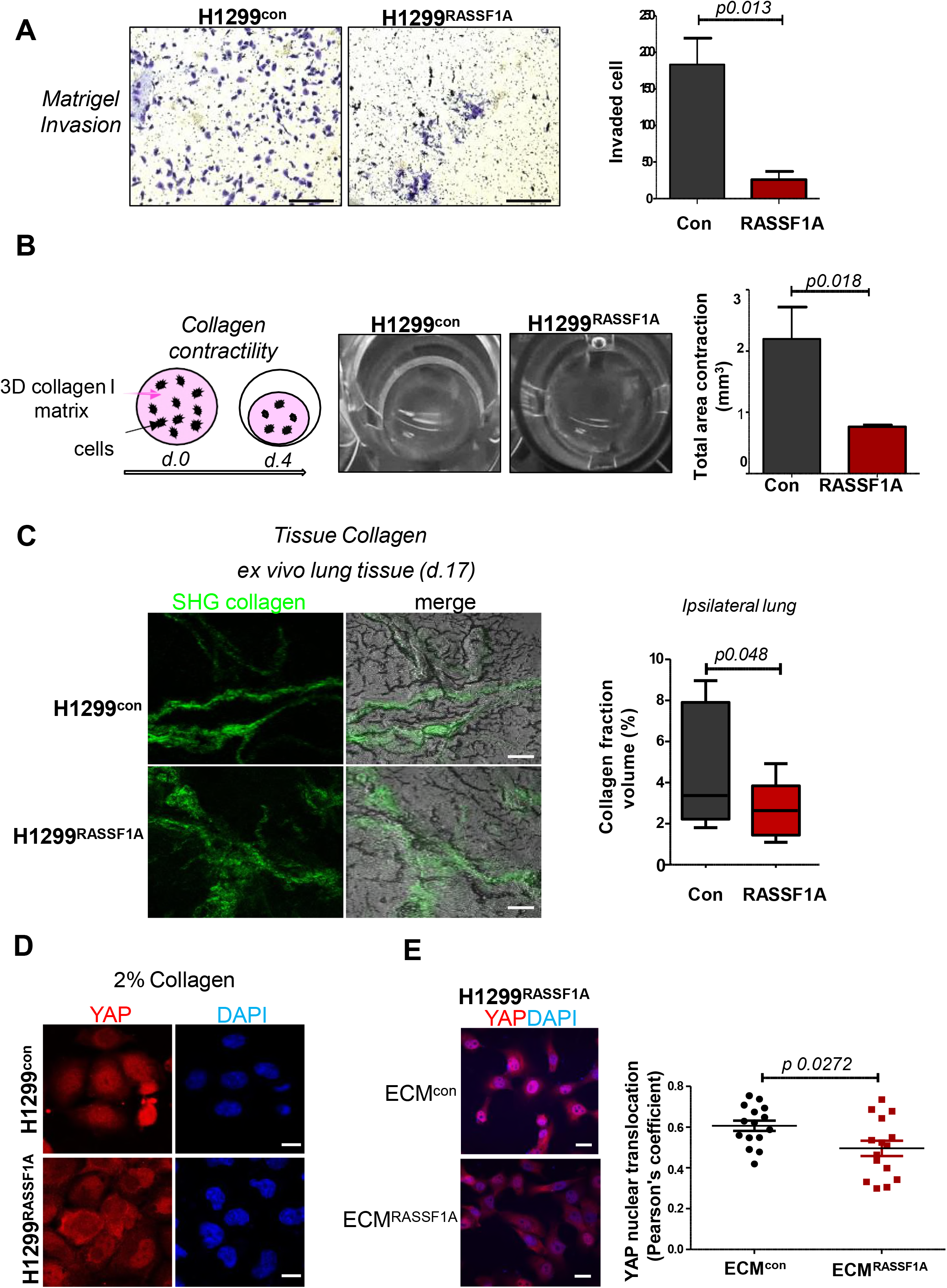
RASSF1A alters invasion and mechanical properties of ECM. **A.** Boyden chamber invasion. Representative images and quantification of invaded cells through the three-dimensional Matrigel matrix, 24 hours, (*n=3*). Scale bars *100μm*. **B.** *Upper:* Cartoon of remodelling assay. *Bottom:* Representative bright-field images showing the effect of H1299 cells on collagen gel contraction after cells were seeded into 3D collagen type I matrices and quantified (*right*) at day 4 (*n=3). P values were determined by using 2-tailed Student’s t-test. Error bars represent mean ±S.E.M*. **C.** Second harmonic generation images of collagen I organization in ipsilateral lung tumour tissue. *Right:* Quantification of collagen I fibrils in lung primary tumours generated at day 17. Scale bars *50μm*. Student’s two-tailed t-test shows a significant difference in collagen I production between control and RASSF1A expressing H1299 cells. **D.** Representative images of H1299 cells grown in 2% collagen. Scale bars *10μm*. **E.** Representative images showing the effect of extracted ECM on YAP nuclear translocation in H1299^RASSF1A^ cells plated and grown in ECM from either H1299^contro1^ or H1299^RASSF1A^. YAP nuclear co-localization (Correlation, Pearson’s coefficient). Scale bars *10μm. Statistical analyses were performed using 2-tailed Student’s t-test*. Error bars represent the mean *±S.E.M. (n=3, 300 cells per experiment*)

We next hypothesized that methylation of RASSF1A in H1299^con^ lung cancer cells may be associated with expression of specific ECM components that are responsible for increased invasion and metastatic dissemination *in vivo*. To identify candidate elements, we performed mass spectrometry analysis of isolated ECM from both H1299^con^ and H1299^RASSF1A^ lung cancer cells. As expected a variety of ECM components were produced by both cells, e.g. collagens, laminins and fibronectin, however, H1299^con^ ECM exclusively contained laminin-beta2 (LAMB2) and prolyl 4-hydroxylase subunit alpha-2 (P4HA2) which contribute to the deposition and stability of ECM (Fig 3A). Interestingly, also exclusive to H1299^con^ ECM was ADAMTSL5 and TPBG (also known as WAIF1 or 5T4; a stem cell receptor that suppresses WNT) which together with P4HA2 all displayed adverse prognostic value for lung adenocarcinoma conversely to RASSF1A expression (Fig 3B). ECM components specifically identified from H1299^RASSF1A^; MRX7A, SDF2, TFPI2 and TIMP1 also had expression profiles that significantly correlated with a good prognosis in lung adenocarcinoma or breast cancer, in keeping with clinical observations for RASSF1A (Fig 3B and Fig EV1D). Interestingly, we showed that high expression of P4HA2 displayed worse prognoses with widespread clinical significance across a range of solid malignancies (Fig EV2A).

**Figure 3.**
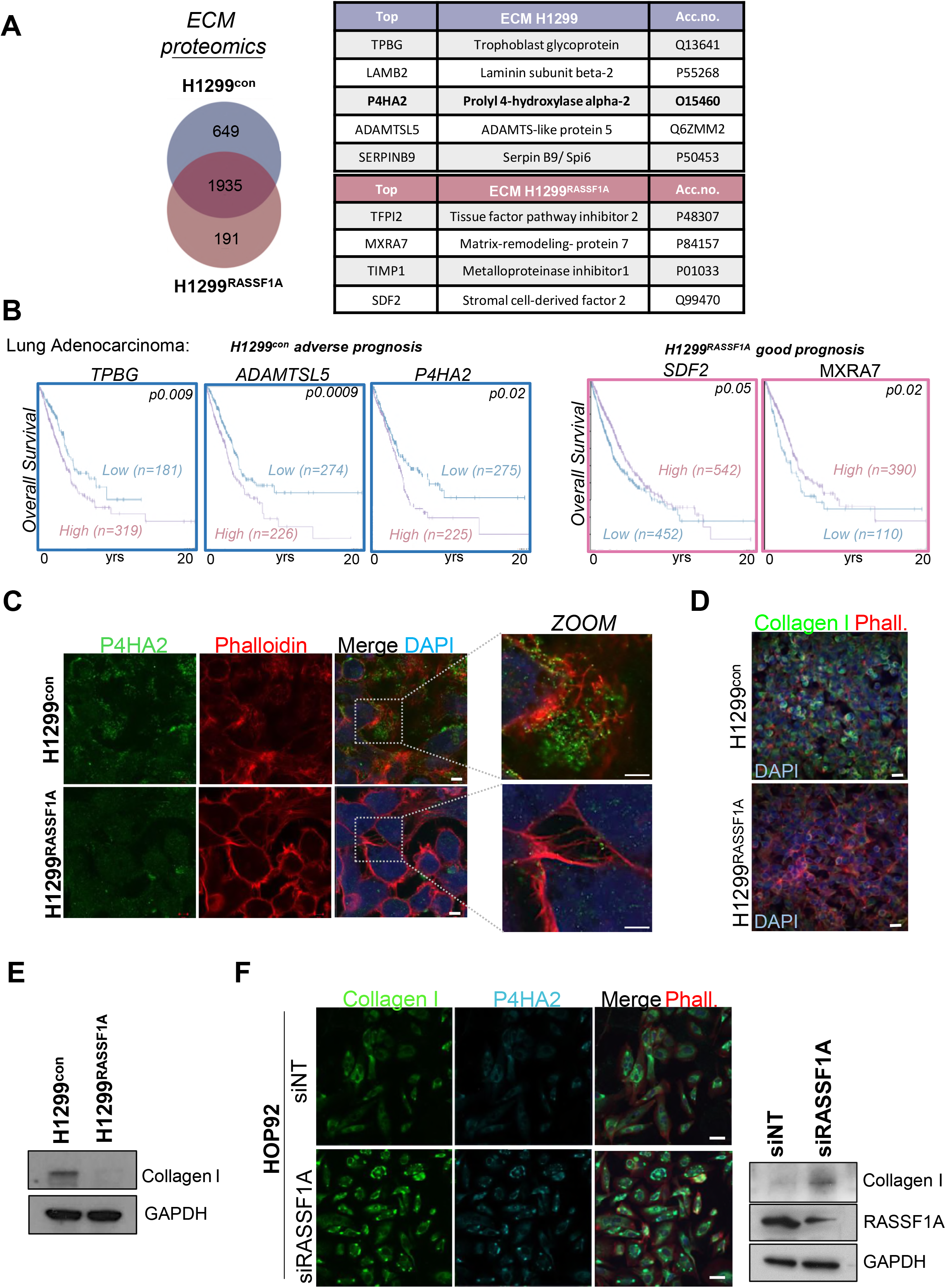
H1299 lung adenocarcinoma cells produce ECM modifying P4HA2. **A.** Mass-spectrometry analysis of proteins purified from extracellular matrix with summary of results. A qualitative analysis of proteins identified was performed using a Venn diagram and only proteins identified with two or more peptides were taken into consideration. The resulting list of proteins was restricted to ECM proteins (*results were provided from triplicates*). **B.** Kaplan Meier plots depicting prognosis in TCGA_LUAD (lung adenocarcinoma) patients with high and low expression of ECM proteins revealed by proteomics study. *The P values were derived from a log-rank test*. **C.** Representative immunofluorescence images of P4HA2 intra and extracellular localization in H1299^con^and H1299RASSF1A cells. Scale bars *5μm, ZOOM* scale bars *2μm*. **D.** Representative immunofluorescence images of expression of Collagen I, co-stained with Phalloidin Alexa-568 in H1299 cells stably expressing either empty lentiviral vector or RASSF1A cultivated on 2D. Scale bars *50μm*. **E.** Western blot analyses of Collagen I expressed in H1299^control^ and H1299^RASSF1A^ cells. **F.** Immunofluorescence staining and western blot analyses of collagen I and P4HA2 in HOP92 cells upon RASSF1A restriction using siRNA. Scale bars *20μm*.

We next confirmed the proteomic data by immunofluorescence staining and did not observe expression of P4HA2 in extracellular space in RASSF1A expressing cells compared to control H1299 cells where RASSF1A is methylated (Fig 3C and Movie EV3, 4). P4HA2 catalyzes the formation of 4-hydroxyproline in collagens and it is essential for proper three-dimensional folding of newly synthesized procollagen fibers (Myllyharju, 2003, 2008). Absence of P4HA2 results in reduced collagen deposition and severe ECM abnormalities (Myllyharju & Kivirikko, 2004). In agreement with P4HA2 expression, H1299^con^ cells display higher levels of stable collagen I compared to H1299^RASSF1A^ cells (Fig 3D, E). To test if RASSF1A is responsible for collagen I switch, we selected a lung adenocarcinoma HOP92 and human cervical HeLa cancer cells that retain endogenous RASSF1A expression and find that these cells displayed low levels of P4HA2 and collagen I, both of which could be upregulated upon silencing of RASSF1A expression with siRNAs (Fig 3F and Fig EV2B).

### P4HA2 expression associates with organized ECM and stiffness *in vitro* and *in vivo*

To investigate if methylation of RASSF1A is associated with the quality of collagen deposition we embedded 3D-spheroids into existing ECM to examine collagen structure by second harmonic generation microscopy (SHG). We found local production of collagen around H1299^con^ spheroids with long, organized fibers radiating from 3-D aggregates which were not visible around H1299^RASSF1A^ spheroids and where appeared as a short, disorganized fibers (Fig 4A). Intriguingly, collagen fibers produced by H1299^con^ cells seem to serve as tracks for invading cells and initiate spread from the central spheroid (Fig 4A, red arrows). P4HA2 is known to have a major effect on physical properties of tumor-associated ECM, which in turn leads to increased stiffness during cancer progression (Provenzano *et al*, 2006; Levental *et al*, 2009). To assess whether our observations correlated *in vivo*, we determined P4HA2 levels in primary tumors by immunohistochemistry (IHC) and, as expected, controls expressed high levels (peri-nuclear due to processing of collagen in the ER; Human Protein Atlas), and reduced staining was evident in H1299^RASSF1A^ tumors (Fig 4B). Correspondingly, SHG microscopy revealed that compared to controls, H1299^RASSF1A^ primary tumors showed significantly lower level of collagen fibers and remaining signal indicated disperse organization with no unifying pattern (Fig 4C), suggesting the reduced levels of P4HA2 affected collagen deposition and may have either prevented metastasis emanating from H1299^RASSF1A^ tumors or restricted their ability to colonize the contralateral lung. Interestingly, H1299^con^ micrometastases mirrored similar collagen deposition surrounding tumorous tissue (Fig 4C, lower) as we observed in 3D spheroids *in vitro* (Fig 4A). We also observed that pre-metastatic control mice (day17) displayed more collagen with organized deposition in the ipsilateral lung (Fig.2C) but not in the contralateral site destined to succumb to metastasis (Fig EV2C). Thus, the variation in collagen level and its organization of the primary tumors is likely to be associated with dissemination rather than colonization. To evaluate how these collagen expression and organization influence the mechanical properties of tumor microenvironment, we next employed atomic force microscopy (AFM) analysis to determine nanoscale variations in stiffness. Consistent with SHG data, detailed topographic analyses of lung tumor stroma by AFM displayed organization of very compact extracellular network mesh with multiple fibers produced by H1299^con^ *in vitro* and *in vivo*, in contrast to H1299^RASSF1A^ (Fig 5A, B). As a consequence, AFM measurements demonstrate H1299^con^ lung tumors tissue were substantially stiffer (16 x kPa vs 1.1 x kPa, *p*<0.001) compared to tumors generated by H1299^RASSF1A^ cells (Fig 5C, D, bars). As the collagen enrichment in tumor ECM is often associated with appearance of fibrotic tissue, contributing to cancer progression (Liu *et al*, 2010), we found that lung tumors from H1299^con^ cells displayed an extended fibrosis that was not apparent in H1299^RASSF1A^ tumors (Fig 5E).

**Figure 4.**
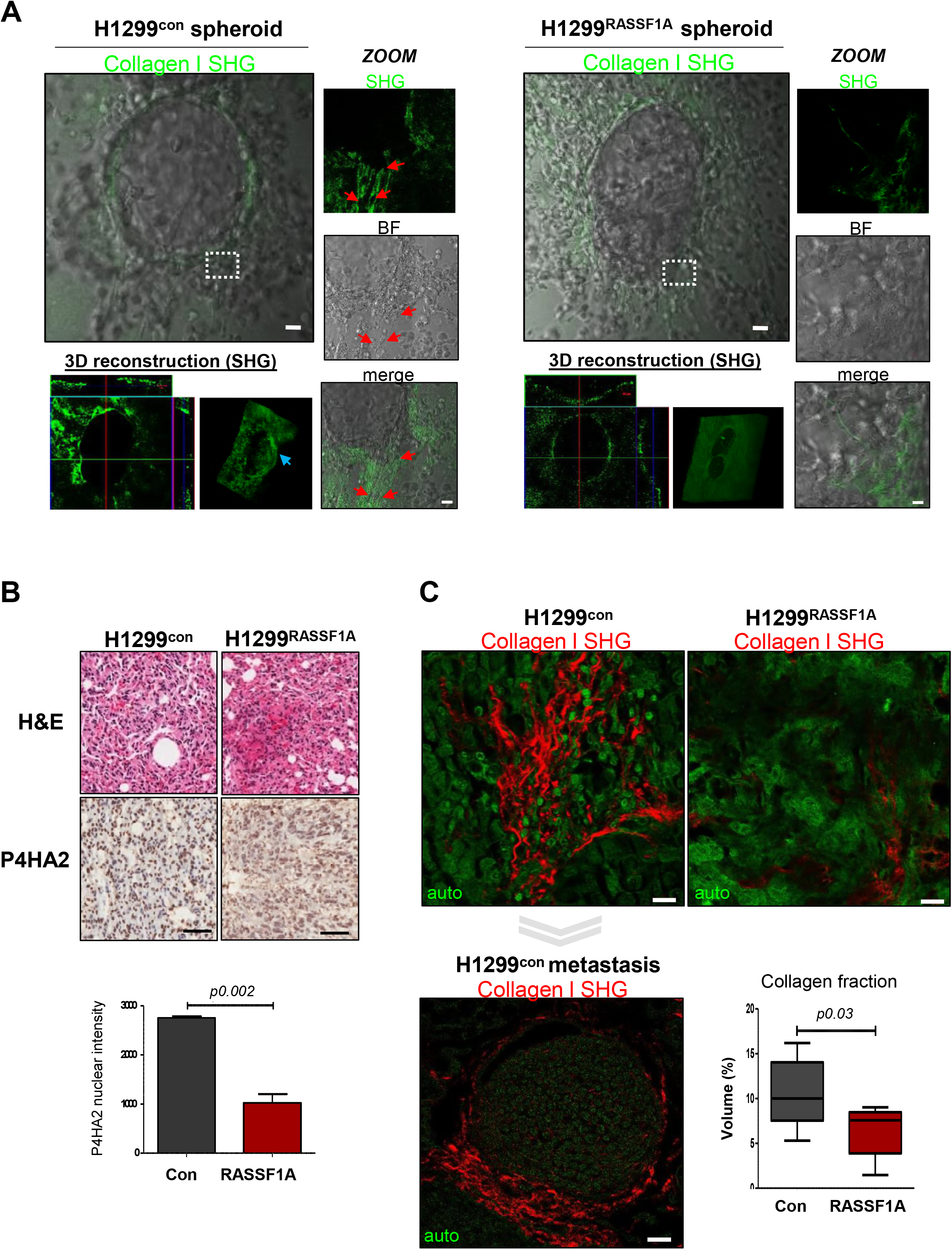
RASSF1A expression associates with P4HA2 and collagen organization *in vitro* and *in vivo*. **A.** Second harmonic generation representative images of Collagen I locally produced by H1299^Control^ or H1299^RASSF1A^ 3D-spheroids, embedded in collagen matrix. Red arrowheads show cancer cell invasion along highly organized, long collagen I fibres compared to disorganized and disperse network produced by H1299^RASSF1A^ spheroids. Scale bars *50μm, ZOOM 20μm*. 3D reconstruction of SHG shows organization of collagen I surrounding spheroids (blue arrowhead). **B.** IHC staining and quantification of P4HA2 expression in lung primary tumours. *P values were determined by 2-tailed Student’s t-test*. Scale bars *100μm*. **C.** SHG images of organization and quantification of collagen I in primary lung tumours at day 30. Scale bars *10μm. Bottom:* SHG representative image of collagen I architecture in micrometastates generated by H1299 control in contralateral lungs. Scale bars *10μm*. Student’s 2-tailed t-test shows a significant difference in collagen I production between control and RASSF1A expressing H1299 generated tumours.

**Figure 5.**
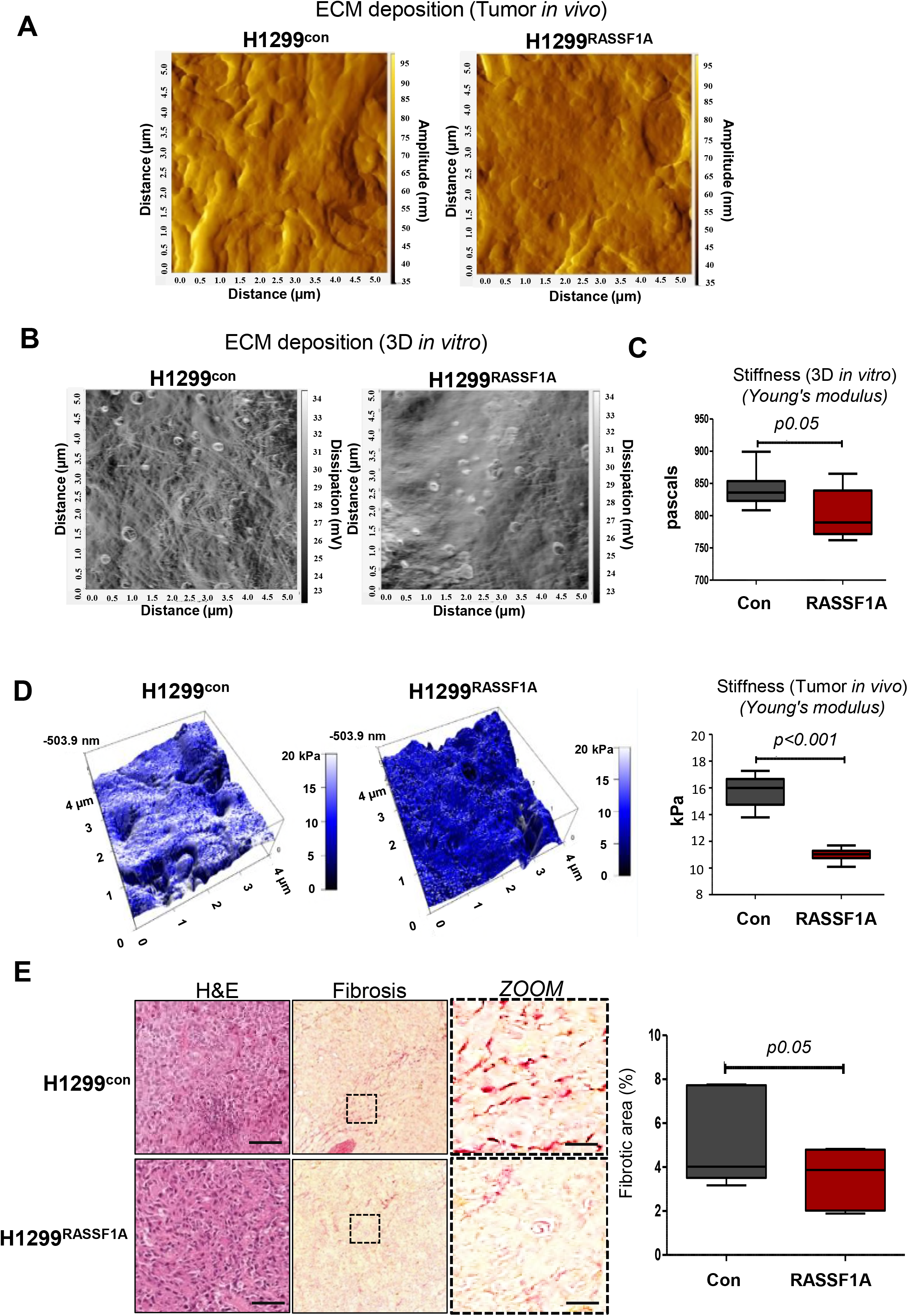
RASSF1A expression is associated with organized ECM and stiffness *in vitro* and *in vivo*. **A.** Atomic force microscopy images showing deposition of extracellular matrix in primary tumours. **B.** Organization of extracellular matrix produced by either H1299^control^ or H1299^RASSF1A^ cells in 3D collagen *in vitro*. **C.** Bar graph representing quantification of stiffness (Young modulus) of ECM generated by H1299 cells *in vitro*, performed by atomic force microscopy. *P values were determined by 2-tailed Student’s t-test*. **D.** *Left:* Representative images of stiffness map in lung primary tumours. *Right:* quantification of stiffness in lung primary tumours. **E.** H&E and Sirius red staining display higher amount of fibrosis found in lung primary tumours implanted with H1299 control lung cancer cells. Scale bars *100μm, zoom 20μm*. Quantification of fibrosis based on Sirius red staining (% of measured area), *Student’s 2-tailed t-test*.

### YAP pathway regulation of P4HA2

The Hippo pathway mediator YAP1 is both a key regulator of P4HA2 levels and has been well documented to be regulated by RASSF1A (Piersma *et al*, 2015; Matallanas *et al*, 2007). Nuclear YAP1 is a readout of transcriptional activation and in agreement with previous data and the expression of P4HA2, YAP1 is significantly more nuclear in H1299^con^ cells compared to H1299^RASSF1A^ (Fig 2D). YAP1 is a pleiotropic transcription factor that drives stemness through activation of the pluripotency cassette (OCT4, SOX2 and Nanog). Disruption of Hippo pathway leads to constitutive nuclear levels of YAP that in turn induces activation of lung cancer stem cells (Noto *et al*, 2017). Despite constitutive nuclear YAP1, H1299^con^ cells grown in 2D did not display upregulation of NANOG compared to H1299^RASSF1A^ and levels remain weak in both cell lines (Fig EV3A). To address whether mechanical properties of ECM can induce stemness, we embedded H1299^con^ and H1299^RASSF1A^ three-dimensional spheroids into the 3D collagen matrix. Strikingly, H1299^con^ spheroids display a substantial level of bright Nanog^+ve^ cells, whereas H1299^RASSF1A^ did not presumably due to reduced nuclear YAP (Fig 6A and Fig EV3B). YAP also regulates Hif1α transcription, an additional activator of P4HA2 under hypoxic conditions, but our spheroid formation method does not result in a hypoxic core, nor do we see Hif1-alpha expression (Fig EV3C). We also show that the appearance of the Nanog+^ve^ cells in the spheroids embedded in three-dimensional collagen matrix was dependent on P4HA2, as the P4HA2 inhibitor 1,4-DPCA abolished nuclear staining of NANOG and reduced nuclear YAP (Fig 6B). To determine whether this effect was a property of 3D-spheroids or cells intrinsic, we next embedded single cells in 3-dimensional collagen matrix and subsequently exposed these cells to P4HA2i. As expected, we found that YAP and NANOG levels in single cells are not affected by acute exposure, but surprisingly we did find that the intrinsically high levels of b-catenin in H1299^con^ cells are immediately sensitive to P4HA2i (Fig 6C). 5T4/TPBG is a transmembrane glycoprotein and stem-cell marker known to restrict WNT signaling and in agreement with Figure 6A, WNT signaling levels are higher in H1299^con^ cells. We interpret this data, together with the isolation of 5T4/TPBG in H1299^con^ ECM to suggest that reduced binding of 5T4/TPBG to the ECM upon P4HA2i allows 5T4/TPBG to suppress WNT signaling and destabilize b-catenin. We found that Nanog^+ve^ cells were similarly dependent on YAP as restriction of YAP expression with siRNA also abolished the stem-like cells population (Fig 6D) and P4HA2 expression in H1299^con^ cells (Fig 6E). So far, our model would suggest that RASSF1A expression suppress YAP nuclear localization, thus fail to express P4HA2 and without the increased collagen deposition and stiffness the ECM cannot impede 5T4/TPBG. As YAP itself is mechanosensitive with ability to translocate into the nucleus independently on Hippo signaling, we were curious whether increased collagen stiffness can also induce cancer stemlike programing. Interestingly, we showed that elevated stiffness of ECM induces production of Nanog^+ve^ cells independently on Hippo pathway (Fig 6F and Fig EV3D). Taken together this data suggests that production of P4HA2 and stiffness is required for maintenance of cancer stem-like phenotype in a YAP and WNT dependent manner.

**Figure 6.**
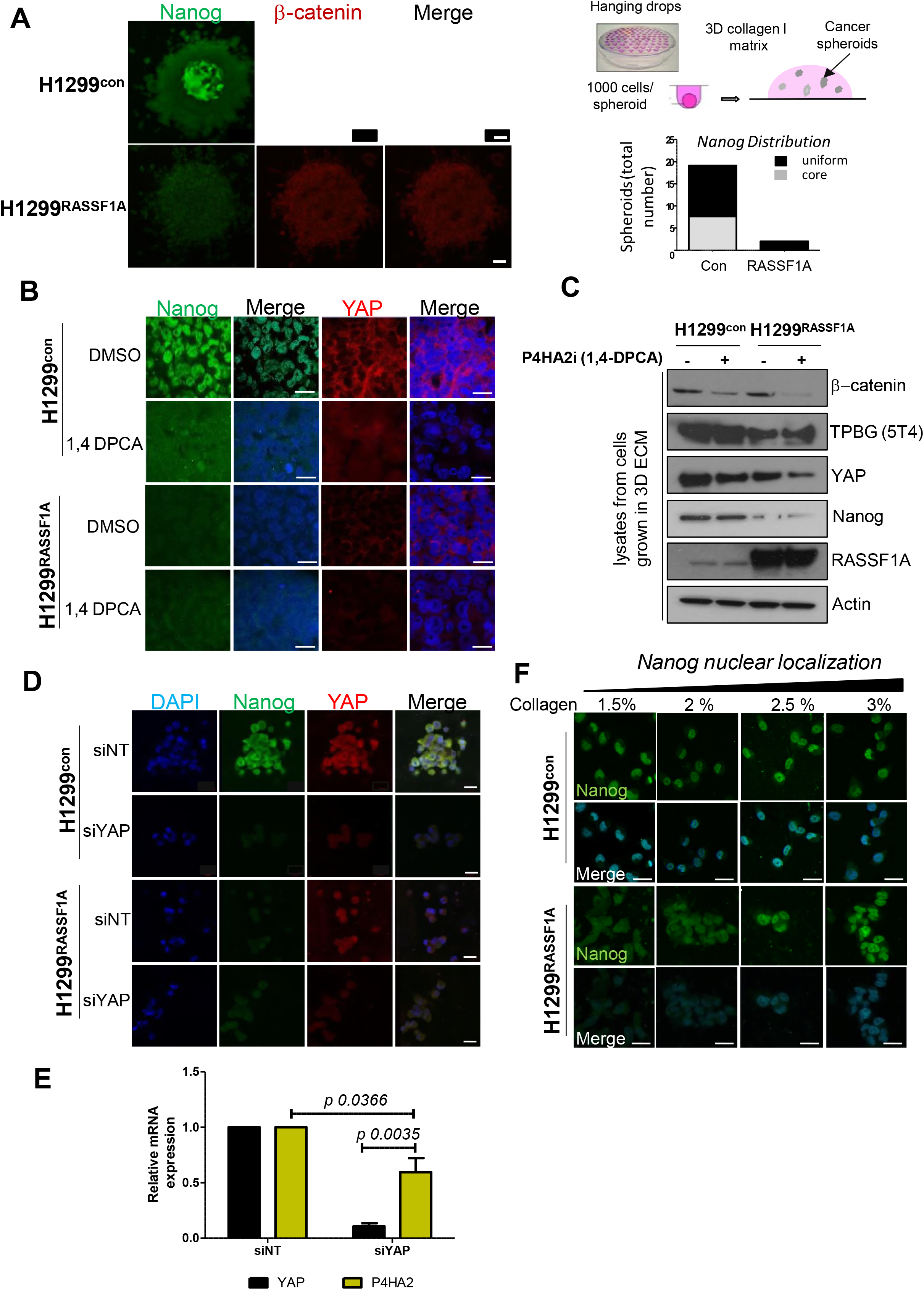
YAP/P4HA2 indirectly regulates WNT via 5T4/TPBG to support NANOG expression. **A.** *Left:* Representative images of immunofluorescence staining for the pluripotency marker Nanog and beta-catenin in 3D-spheroids embedded in collagen matrix. Scale bars *50μm. Right:* Schematic representation of generation of 3D spheroids by hanging drops method. *Lower right:* Quantification of 3D spheroids with Nanog^+ve^ cell distribution either centrally or uniformly across spheroids. **B.** Immunofluorescence staining for Nanog and YAP in 3D spheroids embedded in ECM upon treatment with or without P4HA2 inhibitor. Scale bars *10μm*. **C.** Western blot analyses from H1299 single cells grown in 3D collagen treated or not with P4HA2 inhibitor. **D.** Immunofluorescence images of Nanog and YAP expression with or without YAP knockdown in H1299^control^ and H1299^RASSF1A^ single cells embedded in 3D collagen. Scale bars *20μm* (*n=3*). **E.** Relative mRNA expression of P4HA2 in H1299^contro1^ cells upon YAP silencing with siRNA. *P values were determined by 2-taled Student’s t-test (n=3)*. **F.** Representative immunofluorescence images of Nanog cytoplasmic/nuclear localization (merge image with DAPI) in H1299^control^ or H1299^RASSF1A^ cells growing in 3D matrix with increased stiffness. Scale bars *10μm* (*n=3*).

### Activation of Hippo pathway leads to lung cancer cells differentiation

To address whether the difference in tissue stiffness translated into increased cancer stem-like cells *in vivo*, we stained mouse lung primary tumors and found YAP exclusively nuclear in H1299^con^ tumors compared to more cytoplasmic staining in H1299^RASSF1A^ tumors, and correlated with NANOG expression (Fig 7A). This data supports the model where P4HA2 is required for maintenance of cancer stem-like phenotype and cell pluripotency during early stages of tumor development. Expression pattern of P4HA2 was also highly up-regulated in H1299^con^ metastases but not in the few small metastatic tumors that did appear in mice bearing H1299^RASSF1A^ tumors, suggesting its importance in lung cancer progression (Fig EV4A). We next hypothesized that reduced cancer stem-like cells involvement may imply that tumors expressing RASSF1A are well-differentiated as is observed clinically for *RASSF1* methylation (Grawenda & O’Neill, 2015). To address the question, we evaluated immunohistochemical expression of TTF-1 and Mucin 5B as markers for terminal lung differentiation (Li *et al*, 2012). TTF-1 overexpression in lung tumors tissue has been used as prognostic marker correlated with better prognosis and survival in lung cancer patients (Saad *et al*, 2004; Boggaram, 2009). We first determined levels of Mucin 5B and TTF-1 *in vitro* and found in higher levels in H1299^RASSF1A^ cells compared to controls (Fig EV4B) supporting the idea that the cells were less stemlike and well differentiated. Interestingly, while H1299^con^ spheroids cultured on Matrigel maintained rounded morphology, H1299^RASSF1A^ spheroids collapsed and migrated into branched structures reminiscent of a differentiated epithelium (Fig 7B and Movie EV5). We demonstrated, that high expression levels of mRNA TTF-1 and Mucin 5B correlates with better clinical outcome in patients with lung adenocarcinoma (Fig 7C). To simulate differentiation *in vitro*, we embedded 3D spheroids in three-dimensional collagen matrix and again observed greater expression of TTF1 transcription factor and Mucin 5B in H1299^RASSF1A^ spheroids (Fig 7D). In line with this, primary tumors generated by H1299^RASSF1A^ cells in mice, similarly displayed significantly higher levels of human specific Mucin 5B and TTF-1 compared to H1299^con^ primary tumors (Fig 7E and Fig EV4C). We also observed significant elevation of TTF-1 with its prominent nuclear localization in contralateral metastases generated by H1299^RASSF1A^ cells (Fig EV4D), but failed to observe changes in Mucin 5B expression suggesting this may be specific for primary tumors (Fig EV4D). These data are consistent with a model whereby activation of the Hippo pathway supports differentiation and prevents cancer stem-like cells, ultimately leading to less aggressive forms of lung cancer. Altogether, our results identifies the epigenetic program demonstrating that P4HA2 together with biophysical properties of extracellular matrix drive cancer stem –like programming and metastatic progression in lung cancer and contribute to poor outcome in lung adenocarcinoma patients (Fig 8A).

**Figure 7.**
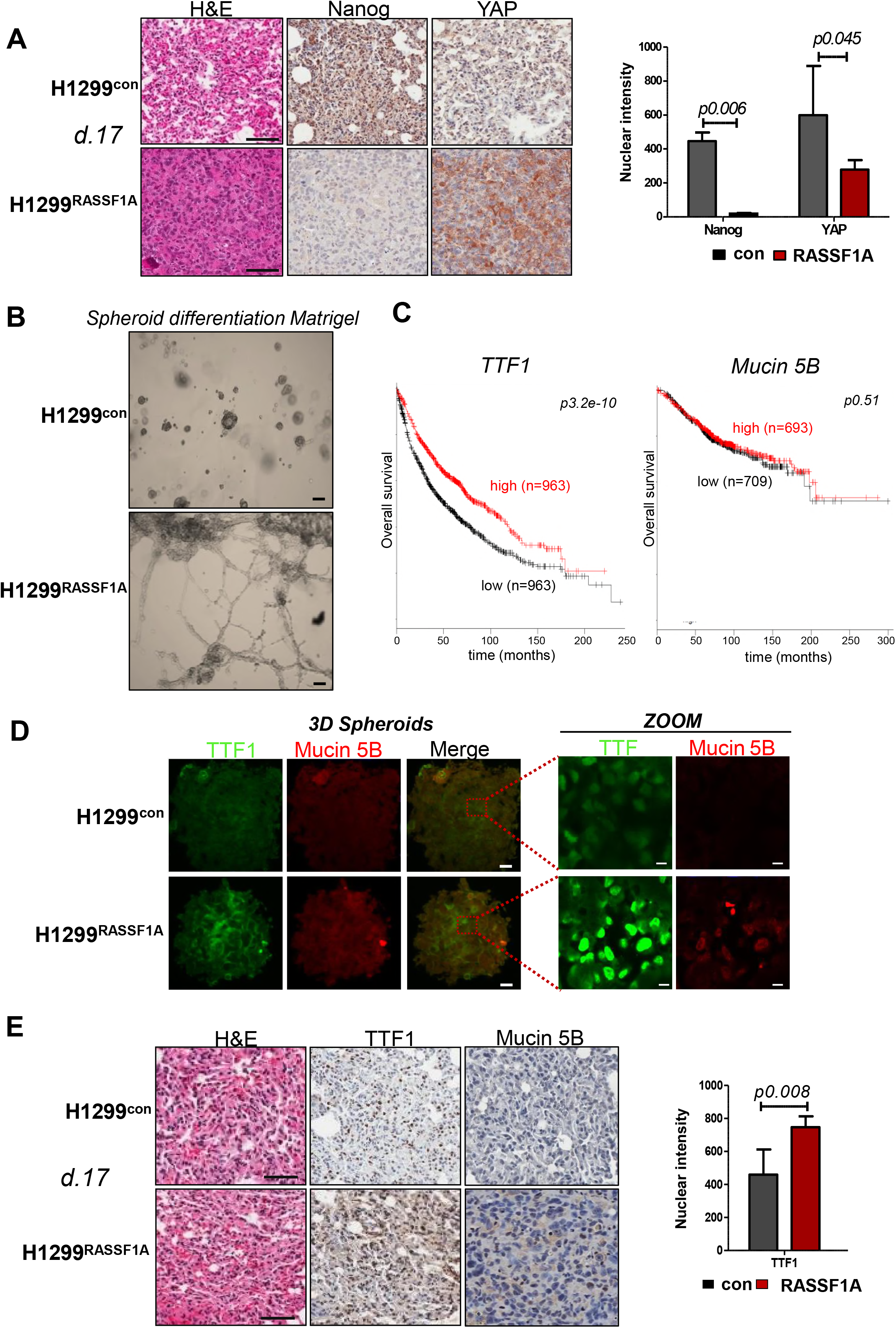
RASSF1A associates with more differentiated lung tumours. **A.** Representative images of H&E and IHC staining for Nanog and YAP in primary lung tumours. Graph bars represent quantification of staining based on strong (3+, 2+) nuclear intensity. *P values were determined by 2-taled Student’s t-test (n=3)*. Scale bars *100μm*. **B.** Differentiation of H1299^control^ and H1299^RASSF1A^ 3D spheroids grown on Matrigel matrix over 24 hours. Scale bars *200μm*. **C.** Kaplan-Meier curves of survival for lung adenocarcinoma patients with high and low expression of TTF-1 and Mucin 5B mRNAs. *The P values were derived from a log-rank test*. **D.** Representative images of immunofluorescence staining for differentiation markers TTF-1 and Mucin 5B expressed in 3D spheroids grown in three-dimensional collagen matrix. Scale bars *50μm, Zoom 10μm*. **E.** Representative images of H&E and IHC staining for lung differentiation markers TTF-1 and Mucin 5B. Quantification of TTF1 expression, based on nuclear intensity in primary tumours on day 17. *P values were determined by 2-taled Student’s t-test*. Scale bars *100μm*.

**Figure 8.**
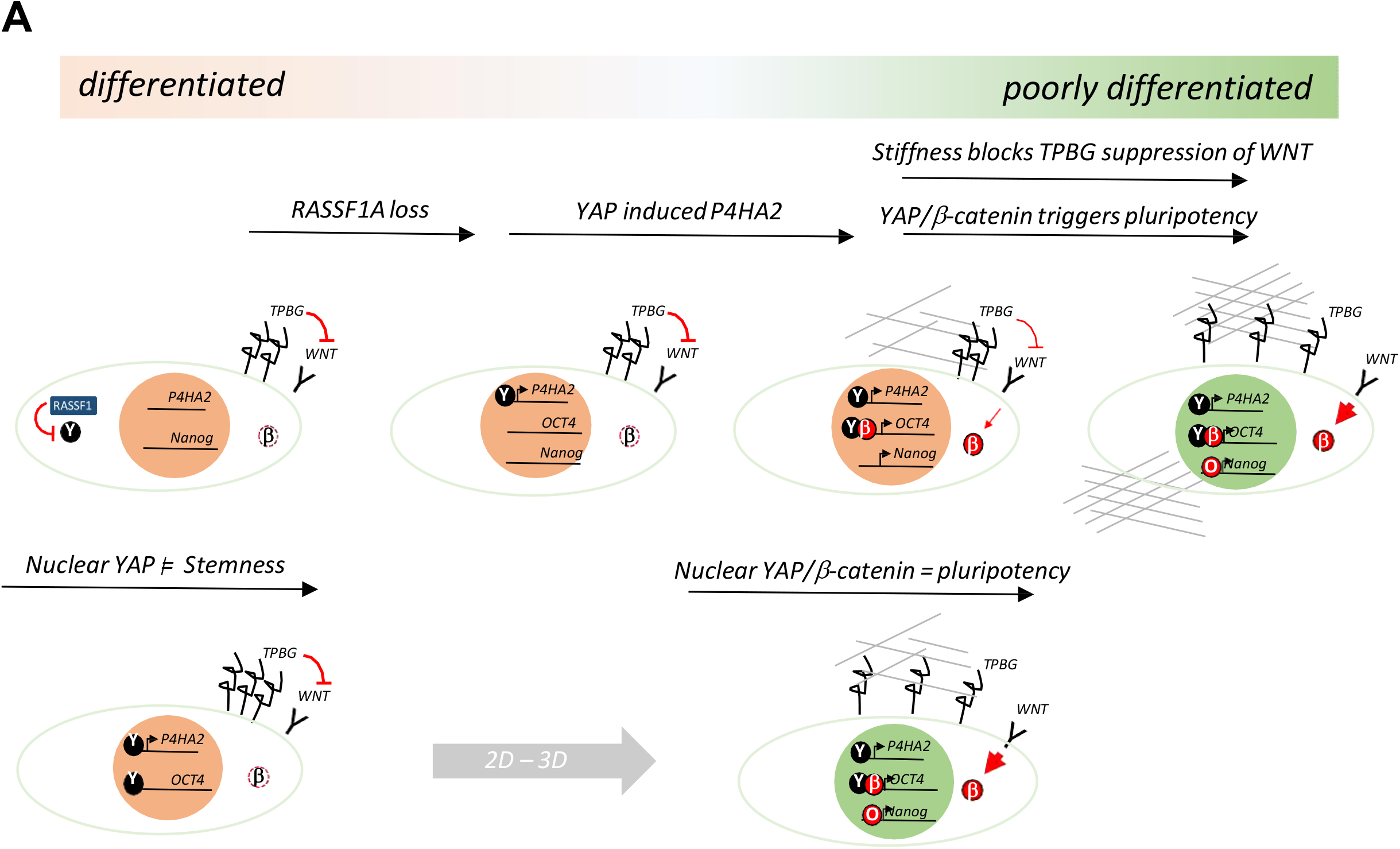
Model for ECM stiffness-promoted stemness in lung adenocarcinoma. **A.** Loss of RASSF1A leads to YAP dependent production of P4HA2 and a stiff ECM. Stiff matrix impedes 5T4/TPBG and prevents inhibition of WNT signaling, allowing beta-catenin stability. Together YAP and beta-catenin activate OCT4 expression and OCT4 commits cells to pluripotency together with Nanog and SOX2. This model implies that nuclear YAP does not activate stemness in the absence of WNT or when repressed, e.g. via 5T4/TPBG. Increased stiffness stabilizes YAP in the nucleus by increasing beta-catenin and promotes OCT4/Nanog and stem cell behaviour.

## Discussion

Lung cancer has a dismal prognosis with only 10% of patients surviving greater than 5 years after diagnosis. As surgery is limited to patients with localized disease, greater confidence in determining which patients are more likely to recur with metastasis is vital to provide appropriate treatment regimens and increase overall survival (Yang *et al*, 2005). RASSF1A is a bone fide tumor suppressor in non-small cell lung cancer, the epigenetic silencing of which correlates with advanced disease, metastatic potential and adverse prognosis (Donninger *et al*, 2007; Dammann *et al*, 2000; Morrissey *et al*, 2001; Dammann *et al*, 2001; Dreijerink *et al*, 2001). However, a precise reason for these clinical associations has been lacking. Here we advance the original epigenetic association of *RASSF1A* levels with poor survival by utilizing mRNA data from The Cancer Genome Atlas (TCGA). We demonstrate for the first time that expression levels can similarly correlate with outcome, but strikingly now show this is specific to adenocarcinomas and no correlation is seen with squamous cell carcinomas of the lung, clarifying from which population the prognostic value is being derived. In line with observations in human tumors, our *in vivo* experiments highlight that RASSF1A only partially restricts tumor formation while the major consequence of loss is increased metastatic spread. Interestingly, metastatic cells are reported to be migrating phenotypically-plastic cells which have the attributes to survive dissemination in the blood stream and colonize distant sites (Van Engeland *et al*, 2002; Folkman, 2002; Woodhouse *et al*, 1997; Fidler, 1999; Chambers *et al*, 2001; Wyckoff *et al*, 2000). We found that lung adenocarcinoma cells lacking RASSF1A expression contain subpopulation of Nanog^+ve^ stem-like cells, and that this population was mostly evident in the context of a tumor or *in vitro* simulated 3D microenvironment.

The primary tumor microenvironment is a heterogeneous and dynamic biological niche which can be composed of distinct populations of cancer cells, stromal fibroblasts and infiltrating immune components (Hanahan & Weinberg, 2011). The mechanical properties also play an important role that can provide a physical barrier and we find that the physical stiffness of the extracellular matrix directly influences the tumor cell heterogeneity. Mechanotransduction allows transcriptional responses to stiffness, in particular the Hippo pathway transcriptional regulator YAP, however how this functions in a 3D-microenvironment or in tumors has not been clear. RASSF1A activates the Hippo pathway to prevent YAP mediated transcription of pluripotency genes such as OCT4, but this requires cooperation from WNT mediated stabilization of beta-catenin for stable promoter binding (Papaspyropoulos *et al*, 2018). OCT4 is a key regulator of Nanog expression and together with SOX2 form the pluripotency cassette activation of which represents true stem cell behavior. Loss of RASSF1A is sufficient to activate YAP transcription that is beta-catenin independent and we find the YAP-target P4HA2 to be elevated in *RASSF1* -methylated H1299 cells compared to RASSF1A expressing controls. P4HA2 is an intracellular enzyme that modifies procollagen I to allow efficient export for contribution to the ECM and can associate with collagen fibers during export. We found that levels of P4HA2 in the extracted ECM were RASSF1A dependent, indicating that collagen processing is highly enriched and concomitantly, we find mature collagen fibers and a much stiffer ECM in control tumors and *in vitro* spheroids.

We show that stiffness of the tumor niche modulates expression of Nanog which, like RASSF1A, predicts tumor progression and poor prognosis, and is important for maintenance of cancer stem cells (Bussolati *et al*, 2008; Ibrahim *et al*, 2012; Zbinden *et al*, 2010). Similarly, P4HA2 not only gives a clear prognostic value for lung adenocarcinoma but across a range of solid tissues where RASSF1A silencing is also predictive and even in lung squamous cell carcinoma where RASSF1A does not – implying potential alternative routes to activating YAP in this setting. Interestingly, we also find an integral membrane protein associated with stem cells and poor prognosis, 5T4/TPBG, in the ECM of H1299 cells. The identification of a membrane protein in the ECM implies a stronger interaction with the ECM in controls as it is maintained during whole cell extraction and the levels of 5T4/TBPG (although higher in controls) do not appear to be RASSF1A dependent. 5T4/TBPG is a repressor of WNT signaling but in control cells WNT is not suppressed and beta-catenin levels are maintained. Inhibition of P4HA2, appears to weaken the ECM sufficiently to allow TPBG to suppress WNT and the reduce beta-catenin stability.

We suggest that mechanism whereby loss of RASSF1A allows YAP mediated expression of P4HA2, collagen deposition and a stiff ECM. The stiffness reinforces nuclear YAP but this is insufficient to drive cancer stem-like cells behavior alone. The ECM binding of 5T4/TPBG restricts its ability to suppress WNT, which then allows beta-catenin and YAP to coordinate activation of the pluripotency cassette via OCT4. The spheroids containing Nanog^+ve^ cancer stem-like cell create highly organized long, very strong collagen branches serve as “tracks” for direct facilitated cell invasion from tumors and encapsulate micrometastases.

Our findings support the idea that the Hippo pathway is important for maintaining differentiation as re-expression of RASSF1A not only suppress cancer stem-like cells but and upregulates lung differentiation markers TTF-1 and Mucin 5B *in vivo* and *in vitro*. TTF-1 (thyroid transcription factor 1), also known as NKX1.2, has been useful marker for differentiating primary lung adenocarcinoma (Moldvay *et al*, 2004) and described as a prognostic marker where moderate or strong TTF-1 expression is associated with better survival (Saad *et al*, 2004; Boggaram, 2009). Overall our data indicates that the tumor microenvironment promotes dedifferentiation and an increase in stem-like behavior, thus targeting P4HA2 in tumor extracellular niche may be an important new strategy of eradicating chemotherapy resistant cancer stem cells during conventional tumor therapy. The work also indicates that RASSF1A levels or promoter methylation can serve as a prognostic biomarker and would be predictive for anti-ECM targeting companion diagnostics.

## Materials and Methods

### Cell culture and drug treatments

Human lung adenocarcinoma cells H1299TetON-pcDNA3 or H1299TetON-RASSF1A under tetracycline promoter were previously described (Van Der Weyden *et al*, 2012; Yee *et al*, 2012). RASSF1A expression was induced after treatment with 1μg/ml doxycycline for 24 hours before experiments. H1299, HOP92 and HeLa cells were cultured in complete DMEM (Gipco) medium, supplemented with 10% foetal bovine serum (Sigma), penicillin/streptomycin (Sigma), and L-glutamine (Sigma) in 5% CO2 humidified atmosphere at 37°C. Cells used in experiments had been passaged fewer than 15 times. For transfections, cells were transfected with siRNA (100nM) using Lipofectamine RNAiMAX (Invitrogen) regarding to manufacturer’s instructions. For generation stable H1299 cell lines, we used human RASSF1A in pBABE system and pBABE-pcDNA3 control. H1299 cells were infected with lentiviruses carrying RASSF1A or control pcDNA3 packaged in 293T cells using GP and VSVG plasmids for 48 hours and then selected with puromycin (10μg/ml) for two weeks. Activity of P4HA2 in cells was inhibited by using of 4 μM, 1, 4-DPCA, a prolyl-4-hydroxylase inhibitor (Cayman Chemical company) for 24 hours.

### Immunofluorescence staining in 3D collagen

H1299 multicellular tumor spheroids were generated by using the hanging-drop method (Foty, 2011). In brief, H1299TetON-pcDNA3 and H1299TetON-RASSF1A cells were induced with 1 μg/ml doxycycline for 24 hours before spheroids formation with or without 1,4-DPCA (Cayman Chemical company). Cells were detached with 2mM EDTA and re-suspended in medium supplemented with methylcellulose (20%, Sigma-Aldrich) and Matrigel matrix (1%, Corning, Growth factor reduced) and incubated as hanging droplets (25 μl) containing 2000 cells for 48 h to generate multicellular aggregates. H1299 spheroids were washed with medium and mixed with rat-tail collagen (Serva, 2.0 mg/mL), 10X PBS, 1M NaOH and complete medium. Spheroids-collagen solution was pipetted as a 100 μl drop-matrix suspension, polymerized at 37°C and replaced with medium. For 3D spheroids staining, collagen-spheroids gels were washed with PBS, fixed with 4% PFA and crosslinked in sodium azide solution overnight. Spheroids were incubated in primary antibodies (Nanog, Cell Signaling; β-catenin, Santa Cruz biotechnology; YAP, Cell Signaling; TTF-1 Thermo Scientific; Mucin5B Santa Cruz biotechnology, HIF-1α, Abcam; dilution 1: 100) in Triton and 10% NGS overnight and after extensive washing, incubated in secondary antibodies (Alexa Fluor-488, Alexa Fluor-594; dilution 1:1000) overnight in 4 °C. Spheroids-collagen gels were washed with PBS and mounted with DAPI. For single cells immunofluorescence staining in three-dimensional collagen, cells were trypsinized, washed in complete medium, counted and (10^5^/ml) cells were mixed with solution containing 1.5 mg/ml; 2mg/ml; 2.5 mg/ml or 3 mg/ml Collagen R (Serva), 10xPBS, 1M NaOH and complete medium. The suspension of cells-gel solution was loaded into 8 wells (Labtech). The gels were polymerized at 37 °C for 30 min and replaced with complete medium. After 48 hours the cells in 3D collagen were stained by protocol described above. Images were captured by using Nikon 10×/0.30 Ph1 objectives.

### Three-dimensional Matrigel migration assay

H1299 multicellular tumour spheroids were generated by hanging drops methods described above. H1299TetON-pcDNA3 and H1299TetON-RASSF1A multicellular aggregates were washed and cultivated on Matrigel matrix (Corning, Growth factor reduced) for 24 hours. Images were monitored at 37 °C using a motorized inverted Nikon Ti microscope (4×/0.10 NA air objective lens) connective to Nikon camera and captured every 30 minutes.

### Boyden chamber invasion assays

RASSF1A expression was induced in H1299 with 1 μg/ul doxycycline 24 hours invasion. After 24 hours cells were trypsinized and (1 × 10^5^) were cultured in serum-free medium in the upper wells (in triplicate) of Transwell Matrigel chambers (Corning) and allowed to invade toward bottom wells supplemented with 10% FBS conditional media. After 24 hours of incubation, invading cells were fixed and stained with Richard-Allan Scientific™ Three-Step Stain-(Thermo Fisher Scientific), photographed, and counted manually using Adobe Photoshop software.

### Quantitative real time PCR analysis

RNA extraction, reverse transcription and qPCR reaction was implemented by using the Ambion^®^ Power SYBR^®^ Green Cells-to-CT^™^ kit following manufacturer’s instructions (Thermo) in a 7500 FAST Real-Time PCR thermocycler with v2.0.5 software (Applied Biosystems). Calculation of mRNA fold change was analysed by using a 2^^^(ΔΔCt) method in relation to the YAP, P4HA2 or GAPDH reference genes. YAP1 sense primer, 5’-TAGCCCTGCGTAGCCAGTTA-3’and antisense primer, 5’-TCATGCTTAGTCCACTGTCTGT-3’; GAPDH sense primer, 5’-TGCACCACCAACTGCTT AGC-3’and antisense primer, 5’-GGCATGGACTGTGGTCATGAG-3’; P4HA2 sense primer, 5’-GCCTGCGCTGGAGGACCTTG-3’ and antisense primer 5’-TGTGCCTGGGTCCAGCCTGT-3’.

### siRNA interference

siRNA interference for silencing RASSF1A was performed using short interfering RNA oligonucleotides GACCUCUGUGGCGACUU (Eurofins MWG). siRNA for silencing YAP1 was performed using short interfering RNA oligonucleotides CUGGUCAGAGAUACUUCUUtt (Eurofins MWG). For non-targeting control siRNA was used sequence UAAGGUAUGAAGAGAUAC (Dharmacon).

### Immunoblotting

For protein analysis on 2D, cells were cultivated on 100-mm dishes and lysed in RIPA buffer as described previously (Palakurthy *et al*, 2009). For protein analyses from 3D H1299 cells expressing either control pcDNA3 or RASSF1A vector were embedded into 3D collagen type I matrix. After 72 hours cells were washed two times with cold PBS and isolated by incubation in collagenase B for 10 minutes at 37°C and lysed in RIPA buffer. Both 2D and 3D lysates were cleared by centrifugation at 15,000 rpm for 20 min and protein concentration was determined by the using the BSA assay, diluted in 2x NUPage sample buffer containing DTT, and incubated at 99°C for 10 min. and lysed in RIPA buffer. The lysates were cleared by centrifugation at 15,000 rpm for 20 min and protein concentration was determined by the using the BSA assay. Proteins (40 μg/lane) were separated on a 10% polyacrylamide gel by SDS-PAGE and transferred to PDVF membrane (Imoobilon-P, Millipore).Non-specific activity was blocked by 1xTBS containing 0.05% tween and 4 % bovine serum albumin (BSA). Membranes were probed with primary antibodies specific for TTF-1 (Thermo Scientific, 1:1,000), Mucin 5B (Santa Cruz biotechnology, 1:500), RASSF1A (Abcam, 1:1000), β-catenin (Santa Cruz biotechnology, 1:500), 5T4 (Abcam, 1:1000), YAP (Cell signalling, 1:1000), Nanog (Abcam, 1:1000), Collagen I (Novusbio, 1:1000) and GAPDH (Cell Signalling, 1:1,000), RASSF1A (eBioscience 1:1000) Membranes were then incubated with the appropriate HRP-conjugated secondary antibodies (Santa Cruz Biotechnology, 1:5000) for 1 hour at room temperature. After extensive washing by TBST, the blots were developed by enhanced chemiluminescence and exposed by using X-Ray films

### Immunohistochemistry

Murine lungs were collected and either followed by cryosectioning (for AFM) or formalin-fixing and paraffin-embedding. Histological sections were deparaffinised, hydrated and exposed to epitope antigen retrieval with ER solution (DAKO, pH6), followed by endogenous peroxidase activity blocking for 5 minutes. After protein blocking (DAKO) for 60 minutes, sections were stained with primary antibodies, using a previously optimized dilution Nanog (Abcam, 1:100), YAP (Cell signalling 1:100), P4HA2 (Santa Cruz Biotechnology, 1:50), TTF-1 (Thermo Scientific, 1:100), Mucin5B (Santa Cruz Biotechnology, 1:50) overnight in 4C. After washing, sections were incubated in secondary antibodies using EnVision detection kit (DAKO) for 30 minutes in room temperatures, rinsed counterstain with hematoxylin and mounted on glass slides. Slides were imaged by using Aperio Scanescope CS slide scanner, and evaluated for and percentage (0%–100%) of nuclear intensity or four-value intensity score (0, none; 1, weak; 2, moderate; and 3, strong) by using ImageScope (Aperio).

### Animal experiments

All animal experiments were performed after local ethical committee review under a project licence issued by the UK Home Office. Lung orthotopic xenografts were generated following a published protocol (Onn *et al*, 2003) with minor modification. In brief, BALB/c nude mice (Charles River Laboratories, U.K) were anaesthetised with 2% isoflurane, and placed in the right lateral decubitus position. 10^6^ of stably transfected either pcDNA3 (control) or Rassf1A expressing H1299 cells in 50 μL of 50% Matrigel (BD Biosciences) were injected into the left lung. Mice were sacrificed on day 17 (n = 4/ group) or day 30 (n = 6/ group), and lungs were collected.

### Immunofluorescence microscopy

Cells were seeded and cultivated on coverslips, fixed in 4% paraformaldehyde for 15 minutes at room temperature and then permeabilized with 0.5% Triton X-100 (Sigma) in PBS for 10 minutes at room temperature. Non-specific binding was blocked with 3% bovine serum albumin (Sigma) in PBS for 30 minutes before incubation with a primary antibody against type I collagen (Novusbio), Nanog (Cell signalling), YAP (Santa Cruz Biotechnology) and P4HA2 (Santa Cruz Biotechnology) for 2 hours at room temperature. Secondary antibodies were applied for 1 hour at room temperature, followed by staining with or without Phalloidin (Life technologies) for 15 minutes and after extensive washing between each step, coverslips were mounted onto microscopy slides with mounting medium containing DAPI. Images were captured by using confocal Nikon 40×/2. objectives. For each condition a minimum of 300 cells were analysed.

### Collagen contraction assay

Collagen gels for contraction assays were prepared by using of 8 parts of Rat tail collagen I (Serva, Germany), 1 part of 10X DMEM, 1 part of 1X DMEM, 1M NaOH (final pH 7.4) and mixed with cells (1 × 10^6^/ml). Gel-cells suspension (triplicates) was pipetted into 24 wells and allowed polymerized in 5% CO2 at 37 °C for at least 30 minutes. The collagen lattices containing cells were detached from the surface of the wells with the sterile needle, replaced with completed medium and allowed to contraction for 4 days. After day 4, the photographs were taken and contraction was calculated as an increase of gel diameters. Comparison of collagen gel contraction was performed by using Student’s unpaired t-test and p-values <0.05 were considered statistically significant.

### Picro-Sirius Red Staining

4 μm paraffin sections were collected on 3-aminopropyltriethoxysilane (AAS) slides and staining was performed in accordance to manufacturer’s protocol (Abcam, ab150681). Briefly, slides were deparaffinised in graded ethanol solutions and then placed in distilled water for 5 minutes. Picro-Sirius Red solution was placed on slides for 60 minutes at RT and then rinsed quickly with Acetic Acid Solution. After staining, slides were passed through graded ethanol solutions, cleared in acetone and mounted with synthetic resin.

### Generation of cell-free extracellular matrix

2 × 10^5^ stably transfected H1299 cells either with pcDNA3 (control) or RASSF1A vector were seeded on coverslips in 12-wells to allow them to produce extracellular matrix for 7 days. Coverslips were washed with PBS and ECM was extracted two times for 5 minutes at 4°C with 0.5% DOC in immunoprecipitation buffer under gentle shaking (Unsöld *et al*, 2001) After discarding cell debris, ECM was washed tree times with PBS to clean ECM of any remaining debris. 5 × 10^4^ H1299 RASSF1A cells were seeded on cell-free extracellular matrix extracted from either control or RASSF1A cells and grown for 3 days. H1299 RASSF1A cells were fixed and stained with YAP antibody.

### Mass-spectrometry

H1299 cells expressing empty pcDNA3 or RASSF1A vector were growing for 2 weeks in presence of 100μg/ml L-ascorbic acid. ECM was extracted by 0.5% DOC (Unsöld *et al*, 2001) and incubated with 0.5 M acetic solution overnight at 4°C. Collected ECM in acetic acid was reduced by 20mM DTT (Sigma), followed by incubation in 30mM iodoacetamide alkylating reagent (Sigma). Proteins from ECM were precipitated via methanol/chloroform extraction and re-suspended in 6M urea in 0.1M Tris pH7.8 by vortexing and sonication. Final protein concentration was measured by BSA. Thirty micrograms of protein was further diluted to bring urea <1M and digested with immobilised trypsin (Fisher Scientific) overnight at 37 °C. Digestion was stopped by acidification of the solution with 1% trifluoroacetic acid (TFA, Fisher Scientific). After digestion, samples were desalted by solid phase using C18+carbon Spin tip (Ltd) and dried down using a Speed Vac. Dried tryptic peptides were re-constituted in 20 μl of 2% acetonitrile-98%H2O and 0.1% TFA and subsequently analysed by nano-LC LS/MS using a Dionex Ultimate 3000 UPLC coupled to a Q Exactive HF mass spectrometer (Thermo Fisher Scientific) (Vaz *et al*, 2016). LC-MS/MS data was searched against the Human UniProt database (November 2015; containing 20274 human sequences) using Mascot data search engine (v2.3). The search was carried out by enabling the Decoy function, whilst selecting trypsin as enzyme (allowing 1 missed cleavage), peptide charge of +2, +3, +4 ions, peptide tolerance of 10 ppm and MS/MS of 0.05 Da; #13C at 1; Carboamidomethyl (C) as fixed modification; and Oxidation (M), Deamidation (NQ) and phosphorylation on (STY) as a variable modification. MASCOT data search results were filtered using ion score cut off at 20 and a false discovery rate (FDR) of 1%. A qualitative analysis of proteins identified was performed using a Venn diagram and only proteins identified with two or more peptides were taken into consideration. The resulting list of proteins was culled to ECM proteins

### Magnetic resonance imaging (MRI)

Mouse MRI was performed on day 30 after H1299 cell implantation using a 4.7 Tesla, 310 mm horizontal bore magnet equipped with a 120 mm bore gradient insert capable of 400 milliTesla/metre (mT/m) in all three axes (Varian Inc, CA). RF transmission and reception was performed with a 30 mm long, 25 mm quadrature birdcage coil (Rapid Biomedical GmbH, Germany). Balanced steady-state free precession (SSFP) scans were acquired (repetition time (TR) = 2.684 ms, echo time (TE) = 1.342 ms, flip angle (FA) = 20°, field of view (FOV) = 48×24×24 mm^3^, matrix = 256×96×96 and RF hard pulse duration = 16 μs). MR images were analysed using ITK-SNAP (Yushkevich *et al*, 2006).

### Mechanical analyses by scanning probe microscopy

5 μm slides from frozen lung tissue were transferred on coverslips and analyzed for stiffness of primary tumors by AFM. For in-vitro three dimensional stiffness analyses, 2 × 10^5^ H1299 cells expressing either pcDNA3 (control) or RASSF1A vector were embedded into 3D rat tail collagen type I matrix and allowed them modified ECM for 5 days in 8 wells (Labtech). 3D collagen gels were transferred onto slides and subjected to mechanical testing by scanning probe microscopy. Scanning probe microscopy was performed on a MFP-3D Atomic Force Microscope (Asylum Research, High Wycombe, UK), with an AC240TS probe (k = 2.0 Nm^−1^, Olympus, Japan). AMFM nanomechanical mapping (Material property measurements using multiple frequency atomic force microscopy, 2008) and loss-tangent imaging (Proksch & Yablon, 2012) were applied, with measurements based on the shift in the probe’s resonant frequencies dependent on the strength of the interaction, or tip-sample contact force (Giessibl, 1997). A tip correction factor was calculated based on the known compressive Young’s modulus of a polycaprolactone calibration sample (300 MPa). Three 5 μm x 5 μm areas were then randomly selected and measured from each material. 512 x 512 pixel maps of height, Young’s modulus and loss tangent were recorded for each image.

### Second harmonic generation using two-photon microscopy

Second harmonic generation signal from collagen were detected upon simultaneous excitation with a 920 nm laser. The signal from collagen was collected using a BP 460/50 filter. The images were acquired using the LSM 7 MP microscope (Carl Zeiss, Jena, Germany) using the Zeiss W Plan-Apochromat 20×/1.0 NA objective lens.

### Statistical analysis

The results from all experiments represent the means ± SD of replicated samples or randomly imaged cells within the field. Numerical values of cell culture and mouse cohort data were analyzed using Student’s t-test for significance in Prism 6 (GraphPad Software Inc.). Unpaired 2-tailed Student’s t-test were used to compare the mean values of 2 groups. The difference was considered as statistical significant of P values < 0.05.

### TCGA survival analysis

The Cancer Genome Atlas (TCGA) project of Genomic Data Commons (GDC) collects and analyses multiple human cancer samples. The TCGA RNA-seq data was mapped using the Ensembl gene id available from TCGA, and the FPKMs (number Fragments per Kilobase of exon per Million reads) for each gene were subsequently used for quantification of expression with a detection threshold of 1 FPKM. Genes were categorized using the same classification as described above. Based on the FPKM value of each gene, patients were classified into two expression groups and the correlation between expression level and patient survival was examined. Genes with a median expression less than FPKM 1 were excluded. The prognosis of each group of patients was examined by Kaplan-Meier survival estimators, and the survival outcomes of the two groups were compared by log-rank tests. Maximally separated Kaplan-Meier plots are presented with log rank P values. Survival analysis was performed using SPSS version 24.0.

## Acknowledgements

We would like to thank Roman Fischer from Discovery Proteomics Facility and Benedikt Kessler from the TDI MS Laboratory, for Mass spectrometry analysis. We would like to acknowledge Prof. Jonathan M. Kurie and Prof. Peter Friedl from M.D. Anderson Cancer Center, Houston, Texas, USA for very valuable comments during writing manuscript. We also thank Sean Smart, Ana Gomes and Danny Allen from imaging core facility for MRI analyses and graphical assistance. We thank to Graham Brown from Microscopy core for his great technical support and Maria Chatzifrangkeskou and Simone Lanfredini for technical assistance. This work was supported by Cancer Research UK A19277.

## Author’s contributions

D.P. designed the research, performed experiments and analysed data. Y.J. and A.R. assisted with animal work, I.V. provided and analysed LC-MS/MS, J.B. performed the fibrotic staining, C.B. performed AFM, E.O.N and D.P. wrote manuscript.

## The paper explained

### Problem

Lung cancer is the leading cause of cancer-related death worldwide with limited improvement in survival over 30 years due to late diagnosis and poor therapeutic options. Epigenetic inactivation of RASSF1A correlates with lung cancer onset, metastasis, poor prognosis and therapeutic responses, however, mechanistic explanation for these clinical associations in human tumours has been lacking.

### Results

Here we demonstrate that RASSF1A methylation in lung adenocarcinomas is a consequence of triggering signals from ECM due to expression of P4HA2 into extracellular matrix, that in turn induces cancer stem-like cells programming and metastatic dissemination *in vivo* and *in vitro*. Re-expression of RASSF1A or inhibition of P4HA2 activity converses these processes and switch into lung cancer differentiation accompanied by better prognosis and clinical survival.

### Impact

Our study demonstrates an importance of tumour microenvironment in epigenetic regulation of cancer stemness and metastatic progression of lung adenocarcinoma and targeting P4HA2 as a new anti-tumour therapy agent.

## Expanded View, Figure legends

**Figure EV 1. RASSF1A suppresses lung adenocarcinoma metastasis. A.** Expression cut off values for percentages of survivals used for high and low expression of RASSF1A protein in lung adenocarcinoma and lung squamous carcinoma patients obtained from TCGA data. *P-values derived from a log-rank test*. **B.** Representative images of lung tumours from mice at day 17 after orthotopic injection with H1299^con^ or H1299^RASSF1A^ cells. Arrows indicate lung primary tumours. **C.** Images of macroscopic appearances of tumour nodules on the lungs at day 30, identified as patchy and whitish areas. **D.** Kaplan Meier plots showing overall survival prognosis in breast cancer patients with high and low expression of TFPI2 and TIMP1 proteins revealed by proteomics study from extracted ECM produced by H1299^RASSF1A^ cancer cells. *The P-values were derived from a log-rank test*.

**Figure EV2. RASSF1A alters properties of ECM by lacking of P4HA2. A.** Clinical outcome and percentage of survival in patients across various cancers shows effect of low versus high expression of mRNA P4HA2. *The P-values were derived from a log-rank test*. **B.** Representative immunofluorescence staining of collagen I and P4HA2 expressed in HELA cells upon RASSF1A restriction using siRNA. Scale bars *50μm*. **C.** Representative second harmonic generation (SHG) images and quantification of collagen I fibrils in contralateral lungs at day 17. Scale bars *50μm. Statistical analyses were performed using 2-tailed Student’s t-test*. Error bars represent S.E.M.

**Figure EV3. Increased mechanical properties of ECM activates stemness in lung adenocarcinoma. A.** Representative immunofluorescence images show Nanog distribution in H1299^control^ and H1299^RASSF1A^ expression cultured in 2D. 2D-scatterplots of pixel intensities show average of Nanog nuclear co-localization (Pearson’s coefficient). Scale bars *10μm*. The quantification of Nanog co-localization represented by Pearson’s coefficient. *P values were determined by 2-tailed Student’s t-test (n=3). Each experiment contains at least 200 cells*. **B.** Representative immunofluorescence images of YAP nuclear or cytoplasmic distribution in H1299^control^ and H1299^RASSF1A^ expressing in 3D spheroids, grown in three-dimensional collagen matrix. Scale bars *100μm, zoom 10μm*. **C.** Representative images of 3D spheroids grown in 3D collagen matrix and stained for Nanog and hypoxia marker HIF-1α. Scale bars *50μm*. **D.** Effect of mechanical properties of extracellular matrix on Nanog nuclear translocation (Pearson’s coefficient) in the H1299^control^ and H1299^RASSF1A^ grown either on 2D or in 3D collagen matrix with increased stiffness. *Each dot represents at least 20 cells. (300 cells per experiment, n=3)*.

**Figure EV4. Expression of RASSF1A is associated with lung tumour differentiation. A.** Representative images of H&E, IHC and quantification of nuclear staining for Nanog, YAP and P4HA2 expressed in contralateral metastases (right lungs) obtained on day 30. Scale bars *100μm*. **B.** Western blot analyses with quantification for lung differentiation markers TTF-1 and Mucin 5B expressed in H1299^control^ and H1299^RASSF1A^ lung cancer cells. **C.** Representative images of H&E and IHC staining for lung differentiation markers TTF-1 and Mucin 5B in primary tumours, obtained on day 30. Scale bars *100μm*. **D.** Representative images of IHC staining for lung differentiation markers Mucin 5B and TTF-1 and quantification of TTF1 expression in (right lungs) contralateral metastases on day 30. Scale bars *100μm. P values were determined by 2-tailed 5tudent’s t-test*.

**Movie EV 1, 2.** Representative MRI movies showing lung primary tumors and lung cancer progression on day 30, generated by orthotopic injection of H1299 control or H1299 RASSF1A expressing cancer cells.

**Movie EV3, 4.** Three-dimensional Z-stack composition. Representative immunofluorescence images of P4HA2 expressed by H1299^control^ cells or H1299^RASSF1A^ cells. Scale bars *5μm*.

**Movie EV5.** Time lapse. H1299^RASSF1A^ 3D spheroids grown on Matrigel matrix undergo collapse of spheroids and differentiate, 24 hours. Scale bars *200μm*.

